# Molecular Dynamics Simulations of the FtsZ mutant G105S

**DOI:** 10.1101/280248

**Authors:** Vidyalakshmi C Muthukumar

## Abstract

In our previous studies we simulated FtsZ monomer and dimer in different nucleotide binding states. In our simulations, we had used the *E.coli* FtsZ homology model including the FtsZ Intrinsically Disordered Region (IDR). Our simulations revealed that FtsZ dynamics involves a key stage in which GTP binds to monomeric FtsZ and opens its nucleotide binding site which in turn favours polymerization. During dimerization, the C-terminal of the top monomer rotates considerably towards the bottom monomer. Such a rotation of the C-terminal domain leads to capture of the nucleotide by its N-terminal domain. In this study we simulate the FtsZ G105S mutant to see if it may have ATPase activity which was reported in a previous study.

## 1. Introduction

In a previous study [1] it was reported that the G105S mutant has an enhanced specificity for ATP ((Adenosine triphosphate) binding and it has temperature-dependent ATPase activity (2- fold increase in initial specific activity at 43 °C compared to 30 °C). *In vitro* FtsZ84 (G105S mutant) has reduced GTPase activity, less than 20% GTP hydrolysis after 80 min [2]. Its ability to crosslink with GTP is reduced significantly, in comparison to the wild-type FtsZ [3]. Temperature dependent increase in GTPase activity is not observed. *In vivo*, FtsZ84 cells do not grow at elevated temperatures [4]. However, at the permissible temperature, Z-rings are formed and cells divide normally but in some cells constriction sites are seen without division, indicating that in some cells FtsZ84 Z-rings are not able to support division even at permissible temperatures. *In vitro*, FtsZ84 has ATPase activity, GTP does not inhibit G105S ATPase activity but the wild-type FtsZ binds to ATP only at very high ATP concentrations (1mM) [1]. ATP is not able to abolish GTP crosslinking with FtsZ [2]. To examine this, we simulated the *E. coli* FtsZ wild-type and the G105S mutant monomers with ATP.

## 2. Methods

### 2.1 G105S monomer and dimer structures

In the G105S mutant (also called FtsZ84), Glycine residue at 105 was replaced by Serine in the *E.coli* FtsZ primary sequence (used in our previous study). Modeller program [5, 6, 7] was used. One model was generated on the basis of a template structure obtained from our previous simulations of the wild-type FtsZ monomer with GDP (coordinates from well equilibrated system, 93.9 ns were used). To generate a PDB file for G105S FtsZ dimer, the protein was aligned with chains A and B of the *Methanocaldococcus jannaschii* FtsZ dimer structure, 1W5A. This alignment was performed using the Multiseq alignment tool in VMD [8].

The starting position of the GDP nucleotide was obtained from the wild-type monomer template referred above. GTP was aligned manually in VMD using the position of GDP for reference. During manual alignment in VMD care was taken to keep a similar orientation of the nucleotide molecule and approximately, the same location in the nucleotide binding site. For ATP, a reference position was not available, therefore, it was simulated from a random initial position close to the nucleotide binding site. For verification, simulations were run for 100 ns and at least 2 independent simulations were performed. Initial position of the nucleotides in simulations are indicated in Fig. 2.1.

**Fig 1.**
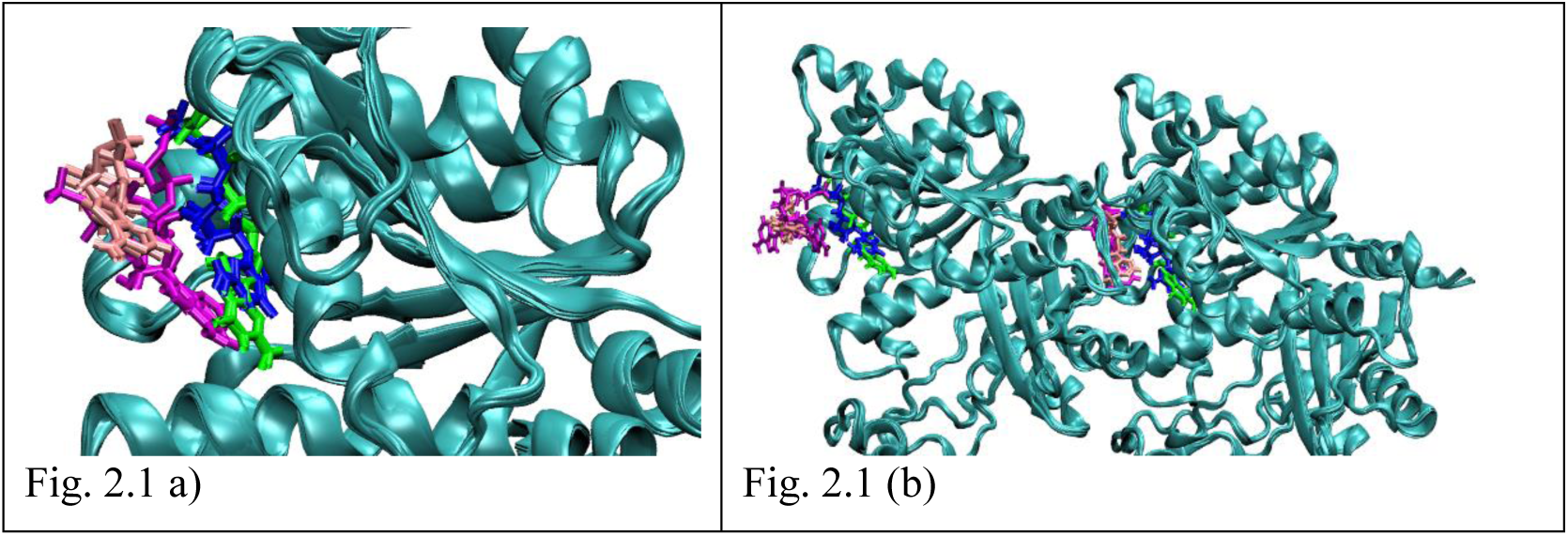
a – b) Position of nucleotides after 100 ps NPT equilibration in (a) monomer simulations (b) dimer simulations. GDP (blue), GTP (green), ATP in wild-type (magenta), ATP in G105S (pink). The wild-type dimer simulation with ATP (Fig. 2.1 (b), magenta) has been discussed in one of our other studies.

### 2.2 Molecular dynamics simulations

Atomistic molecular dynamics simulations were performed using GROMACS version 4.6.5 [9]. Simulations were performed in the AMBER94 force field [10, 11]. AMBER parameters for nucleotides developed by Meagher et al. [12, 13] was used. Protein molecule was centred in the cubic box, at a minimum distance of 1.5 nm from box edges. Solvent (water) molecules were added in the coordinate file. SPC-E (Single Point Charge Extended) water model configuration was used [14]. Mg^2+^ and Cl^-^ ions were added to neutralize the simulation box and at a minimum concentration of up to 10 mM (any of which was higher). Energy minimization was performed using the steepest descent minimization algorithm until the maximum force on any atom in the system was less than 1000.0 kJ/mol/nm. A step size of 0.01 nm was used during energy minimization. A cut-off distance of 1 nm was used for generating the neighbour list and this list was updated at every step. Electrostatic forces were calculated using Particle-Mesh Ewald method (PME) [15]. A cut-off of 1.0 nm was used for calculating electrostatic and Van der Waals forces. Periodic boundary conditions were used. A short 100 ps NVT equilibration was performed. During equilibration, the protein molecule was restrained. Leap-frog integrator MD simulation algorithm [16] was implemented with a time-step of 2 fs. All bonds were constrained by the LINear Constraints Solver (LINCS) constraint-algorithm [17]. Neighbour list was updated every 10 fs. A distance cut-off of 1 nm was used for calculating both electrostatic and van der Waals forces. Electrostatic forces were calculated using PME method. Two groups *i.e.* protein and non-protein (solvent, ligand and ions) were coupled with the modified Berendsen thermostat [18] set at 300 K. The time constant for temperature coupling was set at 0.1 ps. Long range dispersive corrections were applied for both energy and pressure. Coordinates were saved either every 2 ps or every 5 ps (in .xtc compressed trajectory format). A short 100 ps NPT equilibration was performed similar to the NVT equilibration with Parrinello-Rahman barostat [19, 20], with time constant of 2 ps, was applied to maintain pressure at a constant value of 1 bar. 100 ns MD simulation in NPT ensemble was implemented.

## 3. Results and Discussion

### 3.1 MD visualization

The simulations were viewed in VMD. For representation, a frame from the well equilibrated simulation is used. The representative structures are provided in Figure 3.1, taken at 93.9 ns.

**Fig 2.**
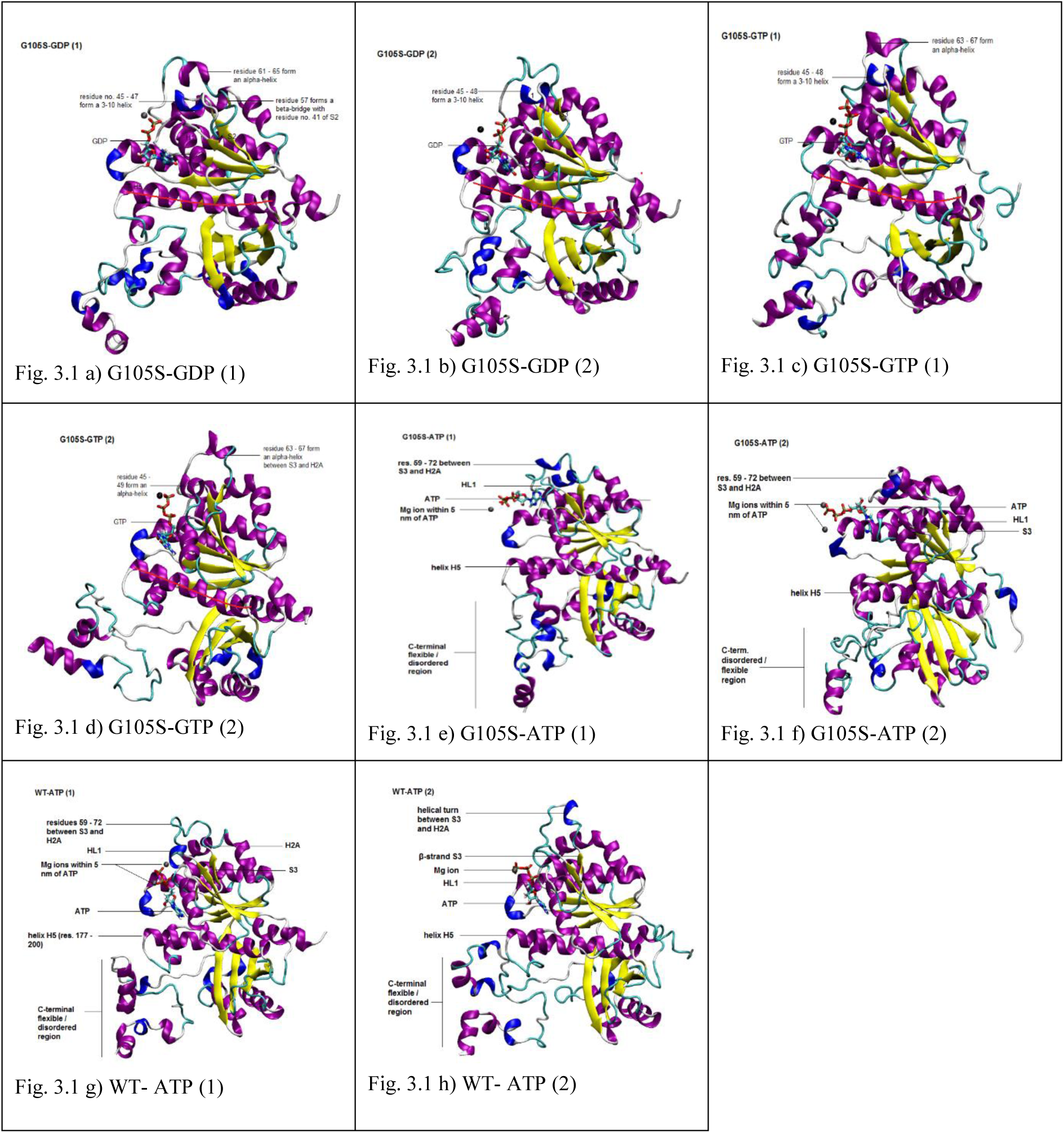
a – d) G105S-mutant structures at 93.9 ns simulation time: (a – b) two G105S-GDP simulations; (c – d) two G105S-GTP simulations; (e – f) two G105S-ATP simulations; (g – h) two WT-ATP simulations. For protein VMD ‘new cartoon’ representation and ‘secondary structure’ colour is used. ‘CPK model’ representation is used for nucleotide and coloured based on atom name; and ‘CPK model’ for Mg^2 +^ ion, coloured in black. The red line along the central helix shows its curvature.

The globular core of the mutant protein is stable in all simulations, with GDP, GTP and ATP. The N-terminal helix and the C-terminal IDR have high flexibility which is consistent with their intrinsically disordered sequence. To our surprise, in the G105S-GDP simulations (Fig. 3.1 (a) and (b)) we observe that the structure of the C-terminal IDR is similar in the two structures. The 3-10 helix and the α-helix are packed very close to the globular core, right below the HL2-H5 loop and close to the C-terminal domain. The nucleotides GDP and GTP occupy stable positions in the nucleotide binding sites in the G105S monomer simulations (Fig. (a) – (d)) and in the G105S dimer simulations (Fig. 3.2 (a) – (d)). The high variations in the position and orientation of GTP which was observed in the simulations of the wild type monomer (reported in our previous study) are not observed in the simulations of the mutant. In the simulations with ATP, it was observed that in the wild-type monomer simulations, the orientation of ATP molecule is such that the adenine points towards the helix H5 and its phosphate tail points outwards (Fig. 3.1 (g) and (h)). The plane of the adenine group is parallel to the H1 helix. This is similar to the orientation of nucleotides GDP or GTP in FtsZ crystal structures. The binding was continuous in both the repeat simulations. In contrast, in the two simulations of G105S monomer with ATP (Fig. 3.1 (e) and (f)), the ATP molecule is observed to have variable/non-specific binding similar to binding of GTP to the FtsZ monomer (observed in our previous study).

**Fig 3.**
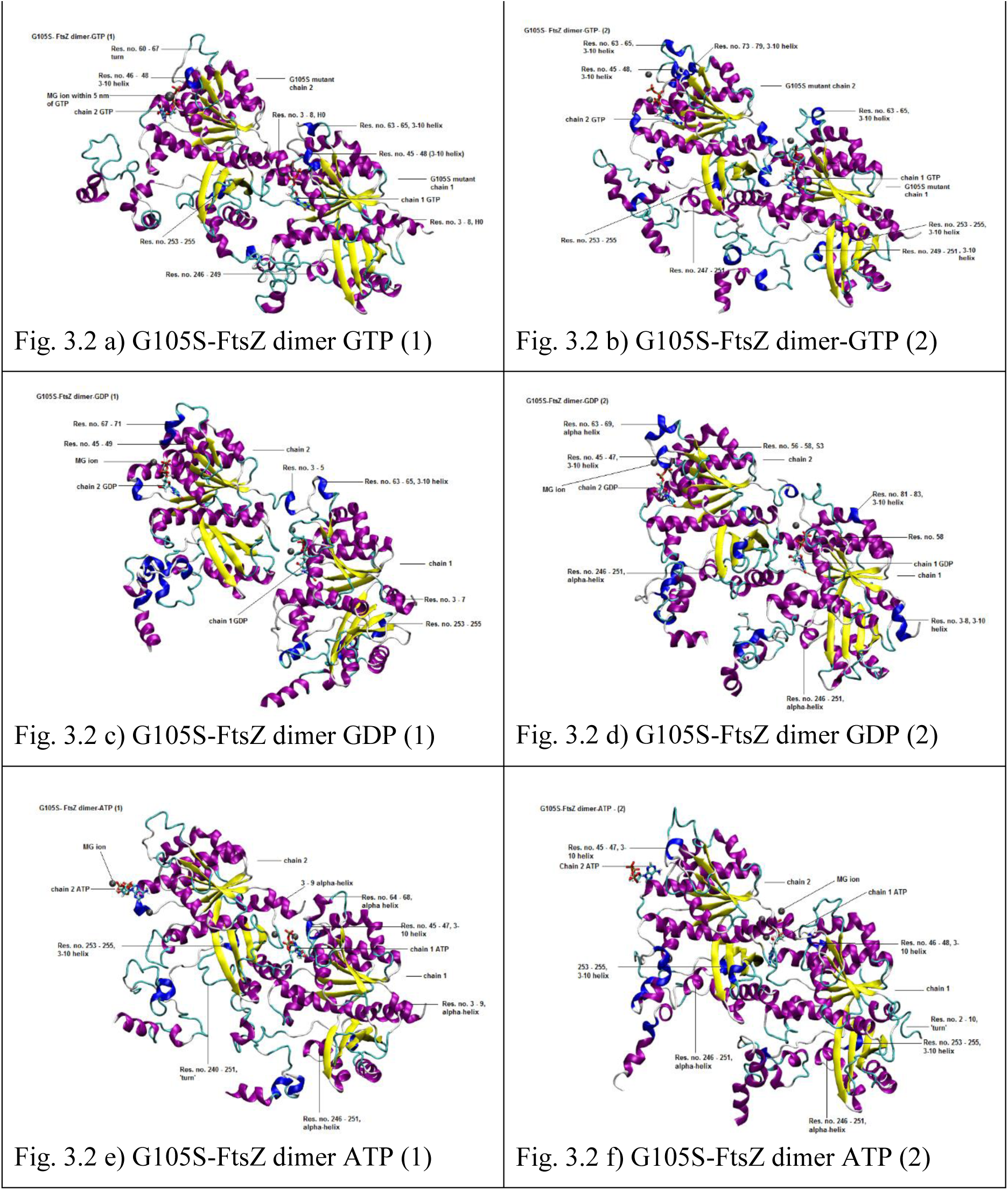
a-f) Structure of the proteins at 93.9 ns of molecular dynamics simulation. The structures of the protein are projected from the same angle in all the figures For protein VMD ‘new cartoon’ representation and ‘secondary structure’ colour is used. ‘CPK model’ representation is used for nucleotide and coloured based on atom name; and ‘CPK model’ for Mg2+ ion, coloured in black. The red line along the central helix shows its curvature. Fig 3.1 a – b) two G105S-FtsZ dimer GTP simulations c – d) two G105S-FtsZ dimer GDP simulations e – f) two G105S-FtsZ dimer ATP simulations.

Variability in conformation of residues 48 – 72 *i.e.* the residues which form the helical loop HL1, the short beta strand S3 and the helical turn/loop between S3 and H2A may be observed in simulations. This is similar to our previous studies and it appears that the region has high flexibility in the mutant simulations also.

Not much change in the curvature of the central helix was seen in any of the simulations. Even in simulations with non-specific binding of ATP, G105S monomer with ATP (Fig. 3.2 (e) and (f)) or G105S dimer with ATP (Fig. 3.2 (e) and (f)), large change in curvature of the central helix could not be seen. The binding of nucleotide GTP does not induce any bending at the central helix, and nucleotide binding appears to be stable. Therefore, it does not appear that the polymerization favouring active state which is observed in the wild-type GTP bound monomer, is formed in the G105S mutant monomer in the presence of GTP, GDP or ATP.

The conformation of residues 245 – 255 *i.e.* the residues between HC2 and SC2 in the C-terminal domain may be found in varying conformations, for e.g. α-helix, 3-10 helix and loop (Fig. 3.1 (c), (g) and (h) respectively).

Binding of ATP to the wild-type monomer does not produce the polymerization favouring active state of the wild-type GTP. Therefore, the polymerization of ATP bound wild-type FtsZ might be quite limited. In our previous study, we had simulated the wild-type dimer with GTP, GDP and ATP. The binding of ATP was found to be non-specific/variable in both the chains of the wild-type dimer in contrast to stable binding observed in the wild-type monomer suggesting that there may be a competition between – binding of the subunits to ATP and dimerization and in this competition dimerization is favoured leading to weak or poor binding with ATP. In contrast, in the wild-type dimer simulations GTP binding facilitates dimerization and the dimer formation in turn leads to stable nucleotide binding.

### 3.2 Root mean square deviation (RMSD)

RMSD calculated for backbone atoms of the residues 12 – 311 are presented in Fig. 3.3 (a – j). The reference structure was the corresponding NPT equilibrated structure. For dimer simulations, protein coordinates are fitted on protein backbone atoms of residues 12 – 311 of both chains. The RMSD profiles are stable in all monomer simulations and for most dimer simulations also. A gradual increase in RMSD is seen in some dimer simulations, for e.g. in G105S-FtsZ dimer-ATP (1) (Fig. 3.3 (i)). Since in the dimer form there may be subunit rearrangement with respect to each other, a higher RMSD can be expected besides RMSD changes due to change in conformation of subunits.

**Fig 4:**
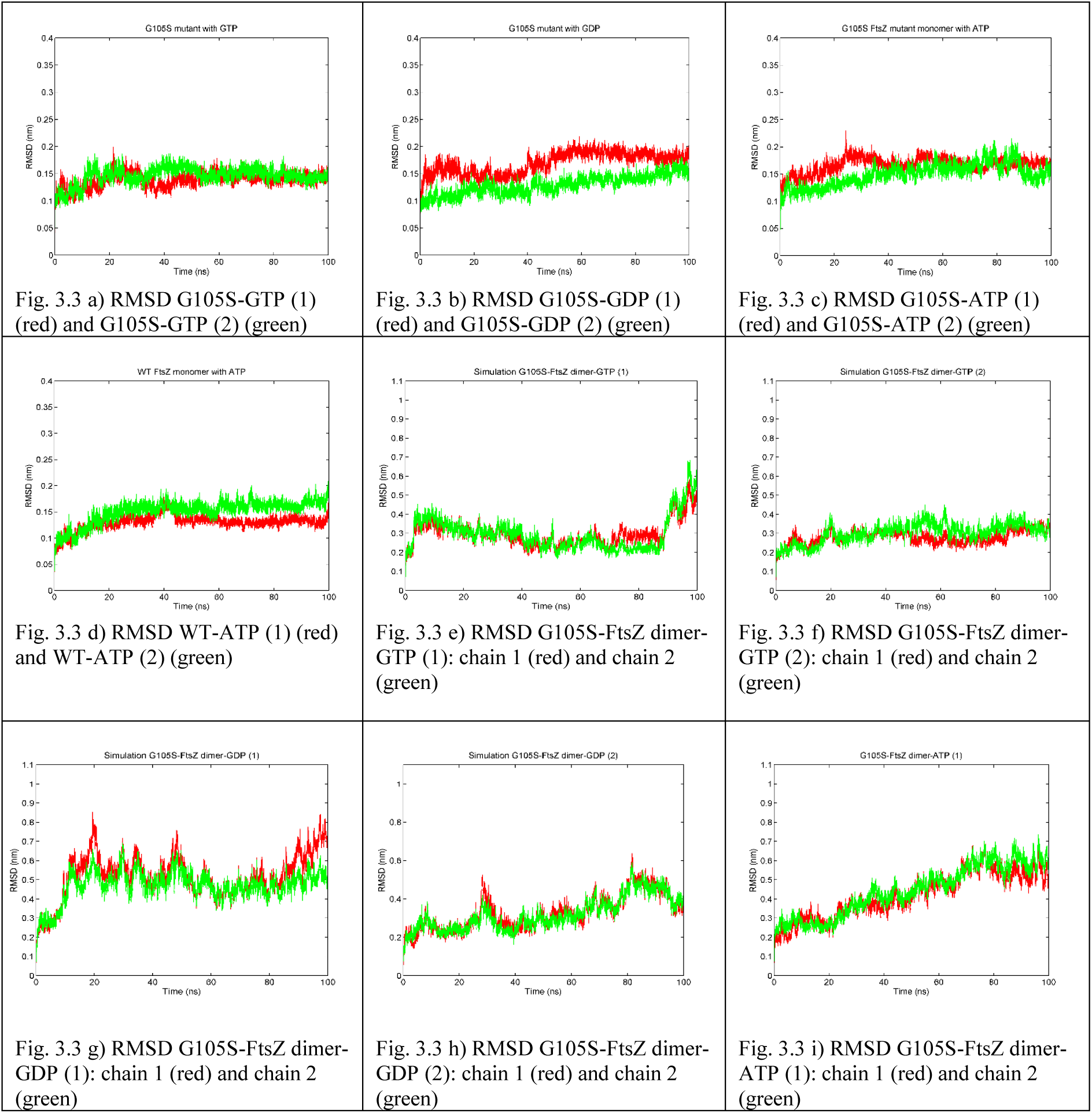
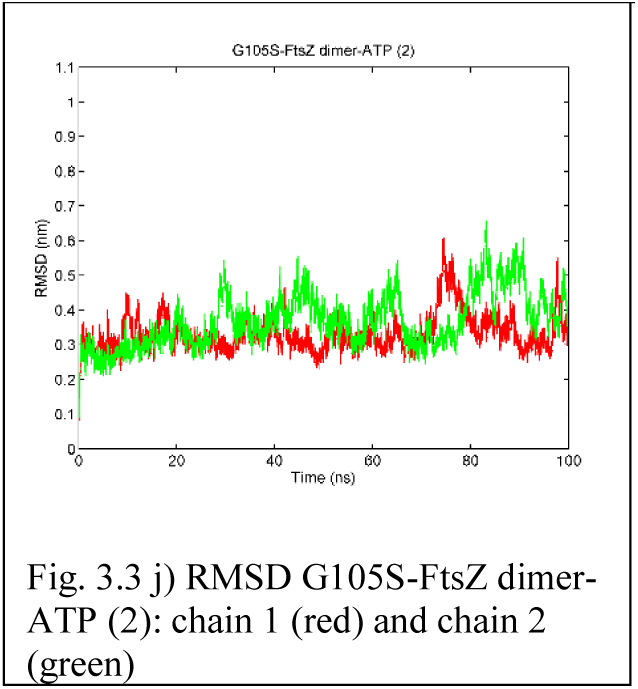
RMSD of backbone atoms of residues 12 – 311 of FtsZ in (a) two simulations of G105S monomer with GTP and (b) two simulations of G105S monomer with GDP (c) two simulations of G105S monomer with ATP (d) two simulations of WT with ATP (e – f) two simulations of the G105S-FtsZ dimer with GTP (g – h) two simulations of the G105S-FtsZ dimer with GDP (i – j) two simulations of the G105S-FtsZ dimer with ATP.

### 3.3 Root Mean Square Fluctuation (RMSF)

RMSF is the standard deviation of atomic positions and is a measure of flexibility of an atom. RMSF of protein backbone atoms (averaged per residue) was calculated. RMSF was calculated from 20 – 100 ns *i.e.* from the well equilibrated system. RMSF for the simulations are presented in Fig. 3.4 (a) – (g). In the G105S mutant monomer simulations, it may be observed that the flexibility of the C-terminal IDR has reduced significantly (Fig. 3.4 (a), (b), (c)). The flexibility of these residues is comparable to or less than what was observed in the wild-type monomer simulations with GTP (SI Fig. 1). The high RMSF of the IDR observed in wild-type monomer simulations with GDP are not observed in any of the G105S monomer simulations. In the wild-type monomer simulation with ATP, the flexibility of the IDR was higher (Fig. 3.4 (d), RMSF in green). Among the dimer simulations, in one of the wild-type dimer simulations with GTP (RMSF G105S-FtsZ dimer-GTP (1), Fig. 3.4 (e)), high RMSF IDR is observed for chain 2 (green).

**Fig 5.**
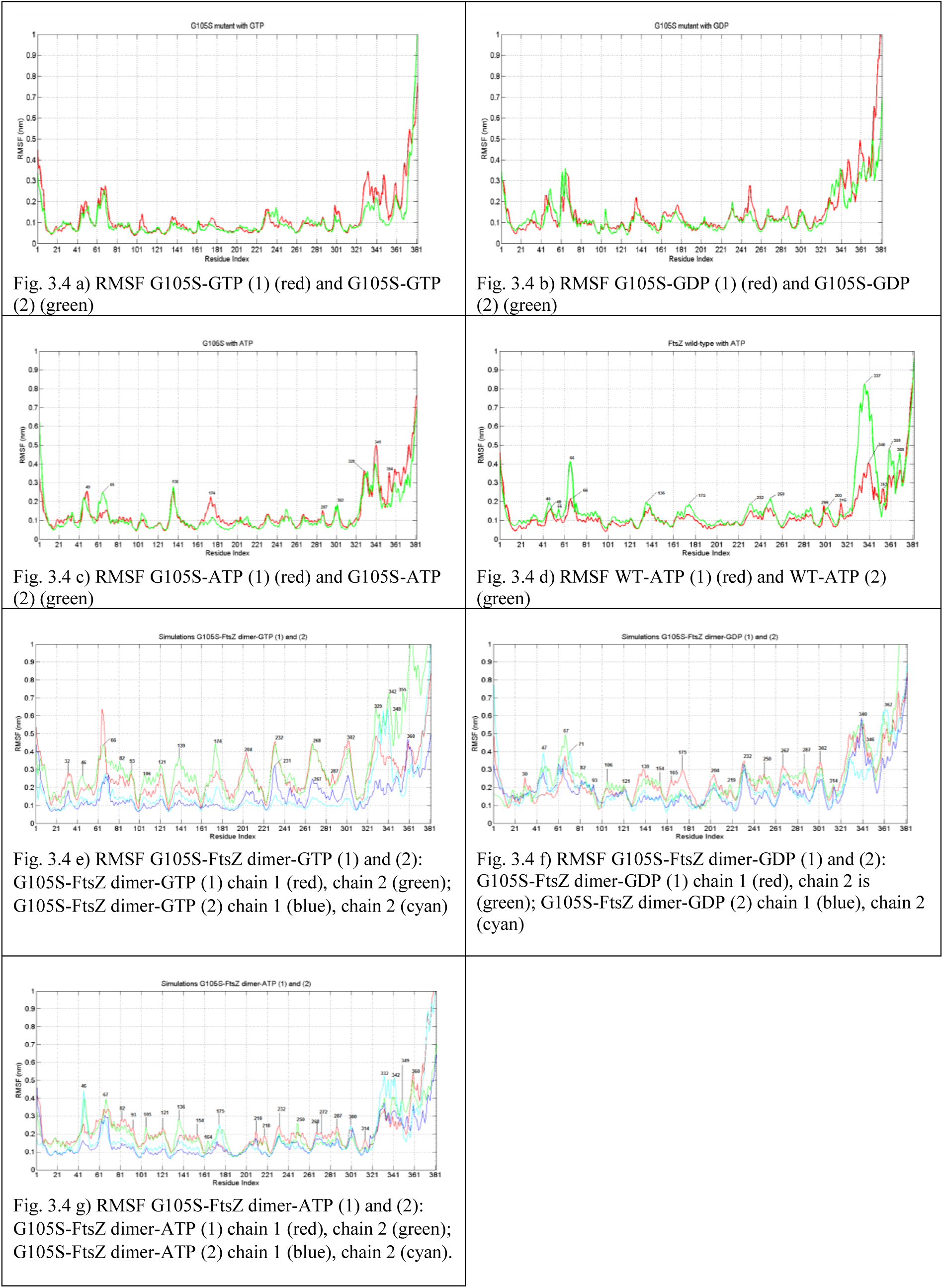
Root mean square fluctuation of protein backbone atoms averaged per residue for (a) two simulations of G105S-GTP and (b) two simulations G105S-GDP (c) two simulations of G105S-ATP (d) two simulations of WT-ATP (e) two simulations of G105S-FtsZ dimer-GTP (f) two simulations of G105S-FtsZ dimer-GDP (g) two simulations of G105S-FtsZ dimer-ATP.

Higher flexibility in the nucleotide binding site residues may be observed (2 peaks between 41 and 81 for S2–HL1 and S3-H2A). The narrow peak between residue no. 161 to 181 which corresponds to helix HL2, loop HL2 – H5 and the N-terminal residues of the central helix which was observed in almost all of the wild type monomer simulations are not observed in the simulations of G105S mutant with GTP. RMSFs for these residues are < 0.2 nm in the two simulations with GDP and just over 0.1 nm in the two simulations with GTP (Fig. 3.4 (a) and (b)). This observation suggests that the flexibility of these residues decreases in the G105S mutant simulations, more so in the presence of GTP. This may be due to stronger hydrogen bonds between the nucleotide and the G105S mutant than in the wild-type protein. In the simulation G105S-ATP (1) maximum RMSF for this peak is just over 0.2 nm (0.2255 nm) andin the simulation G105S-ATP (2) maximum RMSF for this peak is just over 0.1 nm (Fig. 3.4 (c)). RMSFs for the above residues are < 0.2 nm in the two simulations of the wild-type with ATP (Fig. 3.4 (d)).

In Fig. 3.4 (c), it may observed that (the two simulations of the G105S mutant monomer) there is a sharp peak between residues 131 and 141. This peak corresponds to the residue no. 136 which is a Phenylalanine present in the S5-H4 loop. In the two simulation of the G105S mutant, its RMSF is 0.2553 nm and 0.2789 nm (Fig. 3.4 (c)). In the two simulation of the wild-type, its RMSF is 0.1447 nm and 0.1967 nm (Fig. 3.4 (d)). This is in agreement with the non-specific/variable binding of the nucleotide ATP in the G105S simulations, in which ATP can form hydrogen bonds with the outer residues of the nucleotide binding site, therefore increasing the RMSF fluctuation of these residues without actually binding in the nucleotide binding site.

In the simulations of the wild-type dimer (reported in our previous study), RMSF profile showed that dimerization occurs. The N-terminal and the C-terminal domains have high RMSF which rotate about the low RMSF central helix (residues 180 – 200) (SI Fig. 2). We observe similar RMSF profile for simulations, G105S-FtsZ dimer-GTP (1) (Fig. 3.4 (e)) and G105S-FtsZ dimer-GDP (1) and (2) (Fig. 3.4 (f)). In Fig. 3.4 (e), G105S-FtsZ dimer-GTP (2), lower RMSF is observed in the two domains, indicating that dimerization has not occurred. RMSF profile for the G105S dimer simulation with ATP (Fig. 3.4 (g)) does not show high RMSF in the ordered residues which represent domain motions. However, some domain motion in the N-terminal domain appears to have occured in G105S-FtsZ dimer-ATP (1).

### 3.4 Hydrogen Bonds

The residues are denoted by the three letter amino acid code and the residue no. ‘Main’ and ‘Side’ in the residue names indicates that the donor/acceptor atom belongs to the amino acid main chain and to the amino acid side chain respectively. The GTP molecule is denoted as ‘GTP-chain 1’ or ‘GTP-chain2’ referring to the corresponding chains. For calculating hydrogen bonds distance cut-off of 3.5 Å and angle cut-off 25° was used. A hydrogen bond was considered to be significant if occupancy is > 38%.

#### (a) Hydrogen bonds between GDP and the G105S mutant monomer

In the simulation G105S-GDP (1), hydrogen bonds are formed with GLU137-Side, ARG141-Side, PHE134-Main, SER104-Side, PHE134-Side, ASN23-Side. The occupancy of the hydrogen bonds are 91.6%, 89.86%, 70.83%, 68.72% and 43.23%, 37.67% respectively. In the independent repeat simulation, G105S-GDP (2), hydrogen bonds are formed with GLU137-Side, ARG141-Side, PHE134-Main, SER104-Side, ASN23-Side, THR131-Side. The occupancies are 96.36%, 85.08%, 52.95%, 49.12% and 38.55%, 72.41% respectively.

#### (b) Hydrogen bonds between GTP and the G105S mutant monomer

In the simulation G105S-GTP (1), hydrogen bonds are formed with GLU137-Side and ARG141-Side with occupancy 82.47% and 80.27% respectively. In the simulation G105S-GTP (2), hydrogen bonds are formed with GLU137-Side and ARG141-Side and THR107-Side with occupancy 98.92%, 88.28% and 52.92% respectively.

#### (c) Hydrogen bonds between ATP and the G105S mutant monomer

In the simulation G105S-ATP (1), ATP forms hydrogen bonds with ARG141-Side, LYS140-Side. The occupancy of the above hydrogen bonds are 74% and 42% respectively. In the simulation G105S-ATP (2), ATP forms hydrogen bonds with LYS140-Side and LYS139-Side. The occupancy of the above hydrogen bonds are 59% and 51% respectively.

#### (d) Hydrogen bonds between ATP and the wild-type protein

In the simulation WT-ATP (1), ATP forms hydrogen bonds with ARG141-Side, GLU137-Side and ASN23-Side. The occupancy of the above hydrogen bonds are 82%, 55% and 43% respectively.

In the simulation WT-ATP (2), ATP forms hydrogen bonds with GLU137-Side, ARG141-Side and ASN23-Side. The occupancy of the above hydrogen bonds are 96%, 90% and 48% respectively.

It may be noted that in the simulations of the wild-type dimer with GTP, the GTP of top chain (not in the dimer interface) forms hydrogen bonds with Asn23 (reported in our previous study). This raises the possibility that binding with Asn23 might be necessary for secure binding of the nucleotide and perhaps hydrolysis.

#### (e) Hydrogen bonds formed between nucleotides and G105S FtsZ dimer

In the simulation G105S-FtsZ dimer-GTP (1), the GTP at the dimer interface *i.e.* the GTP of the bottom chain, chain 1 (GTP-chain 1) forms hydrogen bonds with THR107-Side, ARG141-Side, THR109-Side, THR109-Main, THR107-Main, ASN23-Side, SER104-Side, GLY108-Main. The occupancies of the above hydrogen bonds are 93%, 92%, 91%, 83%, 74%, 56%, 55% and 47% respectively. The GTP of the top chain, chain 2, (GTP-chain 2) forms hydrogen bonds with GLU137-Side and ARG141-Side. The occupancies are 99% and 88% respectively.

In the simulation G105S-FtsZ dimer-GTP (2), the GTP at the dimer interface *i.e.* the GTP of the bottom chain, chain 1 (GTP-chain 1) forms hydrogen bonds with GLU137-Side, THR107-Side, THR109-Side, THR107-Main, THR109-Main, GLY106-Main, ARG141-Side, THR131-Side, SER104-Main, ASN23-Side, GLY105-Main respectively. The occupancies of the above hydrogen bonds are 98%, 91%, 90%, 69%, 66%, 64%, 61%, 58%, 55%, 53% and 49% respectively. The GTP of chain 2 (GTP-chain 2) forms hydrogen bonds with GLU137-Side and ARG141-Side (occupancies 84% and 73% respectively).

The residues SER104 (which is GLY104 in the wild-type sequence), GLY105, GLY106, THR107 and THR109 belong to the signature sequence in FtsZ. It may be noted that these residues do not bind to GTP in the wild-type dimer simulations. Our results indicate, the in the G105S mutant, the introduction of mutation causes stronger binding with the nucleotide.

In the simulation G105S-FtsZ dimer-GDP (1), the GDP at the dimer interface *i.e.* the GDP of the bottom chain, chain 1 (GDP-chain 1) forms hydrogen bonds with ARG141-Side, GLU137-Side, PHE134-Main, GLY20-Main (occupancies 91.64%, 75.28%, 61.92% and 48.44% respectively). The GDP of the top chain (GDP-chain 2) forms hydrogen bonds with GLU137-Side and ARG141-Side (occupancies 93.01% and 89.01% respectively).

In the simulation G105S-FtsZ dimer-GDP (2), the GDP at the dimer interface *i.e.* the GDP of the bottom chain, chain 1 (GDP-chain 1) forms hydrogen bonds with GLU137-Side, ARG141-Side, GLY108-Main, THR107-Main, ASN23-Side and GLY106-Main (occupancies are 99.0%, 80.30%, 76.81%, 49.63%, 42.39% and 37.53% respectively). The GDP of the top chain (GDP-chain 2) forms hydrogen bonds with GLU137-Side, SER104-Side, THR131-Side, GLY108-Side, ARG141-Side, THR107-Main and THR107-Side (occupancies are 98.25%, 85.79%, 79.18%, 78.80%, 74.56%, 65.96% and 59.85% respectively).

The ATP molecule forms fewer hydrogen bonds than the GTP and GDP discussed above. The hydrogen bonds formed in the two repeat simulations with ATP are described below. In the simulation G105S-FtsZ dimer-ATP (1), the ATP at the dimer interface *i.e.* the ATP of the bottom chain, chain 1 (ATP-chain 1) forms hydrogen bonds with ARG141-Side and LYS140-Side (occupancies are 79.78% and 40.82% respectively). The ATP of the top chain (ATP-chain 2) forms hydrogen bonds with LYS139-Side, ARG141-Side and LYS139-Main (occupancies 73.78%, 71.91% and 68.41% respectively). It forms hydrogen bonds mainly with the residues of the S5-H4 loop of the nucleotide binding site. In the simulation G105S-FtsZ dimer-ATP (2), the ATP at the dimer interface *i.e.* the ATP of the bottom chain, chain 1 (ATP-chain 1) forms hydrogen bonds with LYS140-Side, PHE136-Main and ARG141-Side of chain 1 and ASP211-Side and ASP287-Main of chain 2 C-terminal domain (occupancies are 82.15%,56.93%, 56.68%, 46.57% and 35.21% respectively). The ATP of the top chain (ATP-chain 2) forms hydrogen bonds with LYS140-Side and ARG141-Side occupancies 73.78% and 67.79% respectively).

### 3.5 Principal component analysis

Principal component motion along the first eigen vectors for the monomer and dimer simulations are shown in Fig. 3.5 and 3.6.

**Fig 6.**
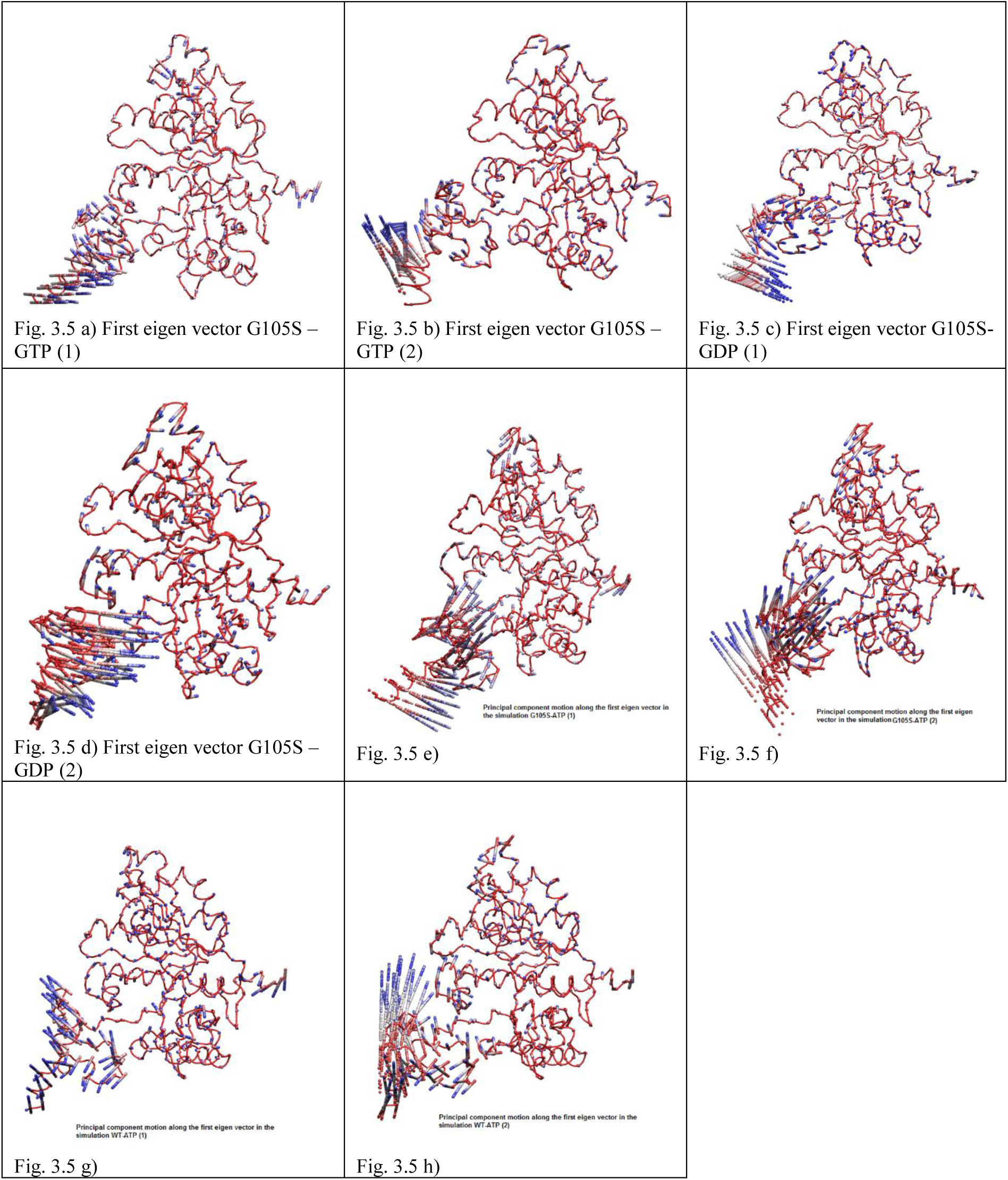
Principal component along the first eigen vector for simulations (a – b) G105S-GTP (1) and (2) (b – c) G105S-GDP (1) and (2) (e – f) G105S-ATP (1) and (2) (g – h) WT-ATP (1) and (2).

**Fig 7.**
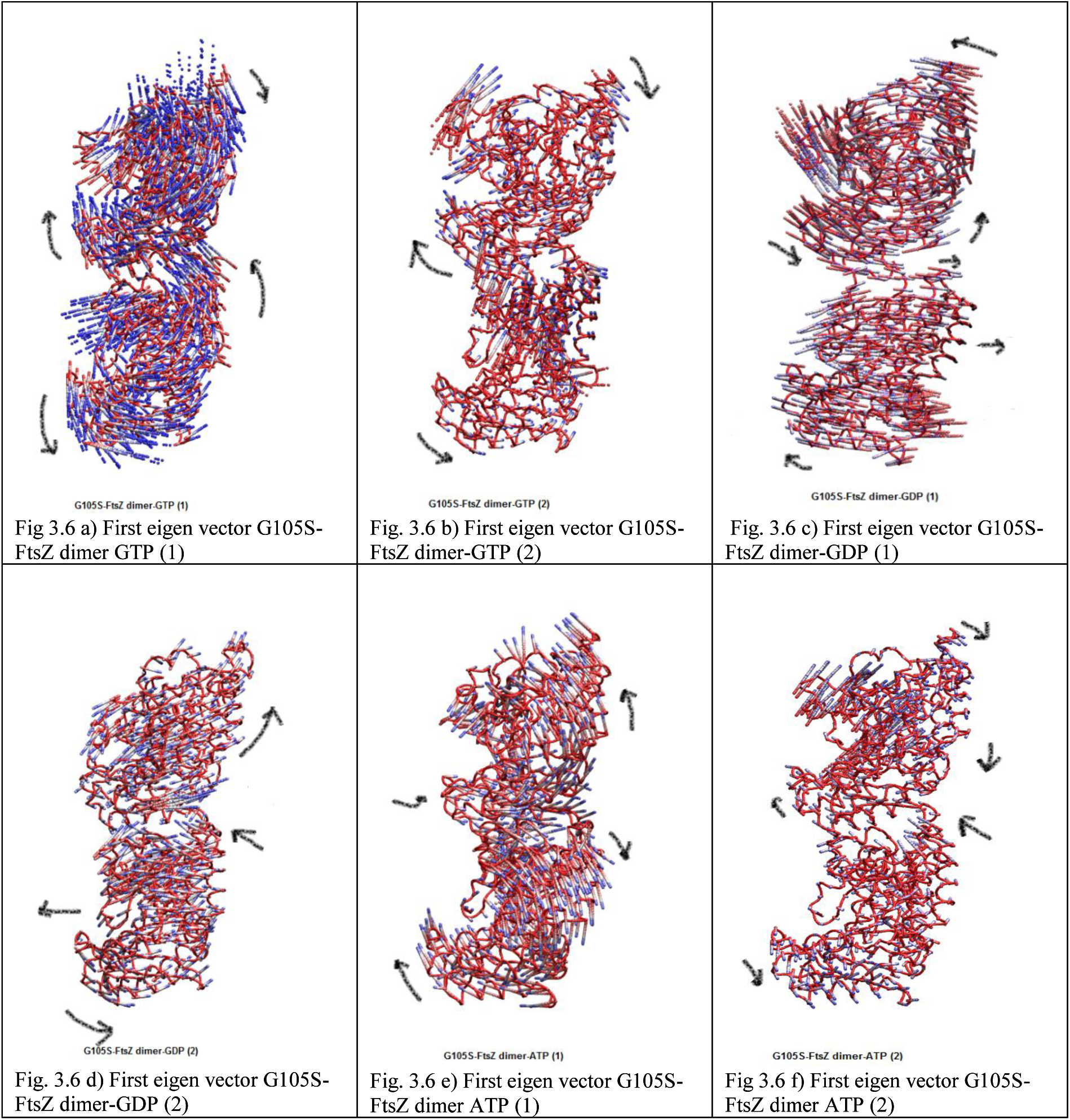
a – f) Principal component along eigen vector 1 for simulations G105S mutant FtsZ dimer with nucleotides (a – b) GTP, (c – d) GDP, (e – f) ATP. PCA analysis was done for Cα atoms. The filtered trajectory along the first eigen vector is represented. The structure of the proteins are projected from the same angle in all states. The ‘tube’ representation in VMD is used to show initial frame (*i.e.* Frame 1, at 5 ps MD simulation). VMD “trajectory” colouring method is used with the option ‘timestep’ which colours atoms according to simulation time (red is initial frame and blue final frame). To show the motion of C-alpha atoms, the ‘CPK’ representation in VMD is used to show the frames from *t* = 0 to *t* = 100 ns with 1000 ps interval. VMD “trajectory” colouring method is used with the option ‘timestep’ which colours atoms according to simulation time (red is initial frame and blue final frame). The images were rendered in VMD. Arrows were drawn to indicate the direction of rotation of N and C-terminal domains or the direction of rotation of the chains.

In Fig. 3.5 (a) and (b), simulation of the G105S monomer with GTP, the displacements in globular core are very low, C-terminal domain motion is not observed. There is no displacement of the S5-H4 loop or in the HL2-H5. Some displacement in the N-terminal helix H0 and the S3-H2A loop is observed. In Fig. 3.5 (a) in addition, displacement of the HL1, S3 in the direction of the nucleotide is observed. The magnitude of motion is similar to the magnitude of motion of the corresponding C-terminal domain residues in the simulations of the wild-type monomer with GDP and less than the magnitude of motion of the corresponding residues in the simulations of the wild-type monomer with GTP (results from our previous study). In Fig. 3.5 (c), simulation G105S-GDP (1), the displacements are similar to the G105S-GTP simulations. In Fig. 3.5 (d), simulation G105S-GDP (2), larger displacements are seen in the N-terminal helix H0, HL1, S3, S3-H2A, S5-H4, HL2-H5. Some rotation of the C-terminal domain is observed. In the simulations G105S-ATP (1) and (2), Fig. 3.5 (e) and (f), larger displacements are seen in the N-terminal helix H0, HL1, S3, S3-H2A, HL2-H5 and some rotation of the C-terminal domain may be seen. In Fig. 3.5 (e) (G105S-ATP (1)), the direction of displacement of the HL2-H5 loop is away from the nucleotide binding site (downwards in HL2 and towards the front in the N-terminal residues of helix H5). In Fig. 3.5 (f) (G105S-ATP (2)), the direction of displacement of the HL2-H5 loop is towards the nucleotide binding site (upwards in HL2 and in the N-terminal residues of helix H5). Displacement in a few S5-H4 loop residues may also be observed (Fig. 3.5 (f)). In Fig. 3.5 (g), WT-ATP (1), displacement are seen in H0, S3-H2A and some C-terminal domain motion. In Fig. 3.5 (h), displacement in H0, S3-H2A and HL2-H5 is observed.

In our previous study we had discussed the rotation of the two chains in the wild-type dimer simulations. The top chain rotation anti-clockwise and the bottom chain rotates clockwise. The C-terminal domain of the top chain is accommodated into the nucleotide binding site of the bottom chain during this rotation. Principal component motion along the first eigen vector for the simulations G105S-FtsZ dimer-GTP (1) and (2) are presented in Fig. 3.6 (a – b). The top chain, chain 2, rotates clockwise and the bottom chain rotates anti-clockwise. This rotation disfavours polymerization since the top C-terminal domain moves away from the dimer interface. This suggests that while the G105S dimer also binds to GTP and is present in a polymer form, the polymer form is similar to the GDP bound form with a higher curvature. Less rotational motion is seen in the bottom chain of G105S-FtsZ dimer-GTP (2) N-terminal domain indicating that polymerization may not be efficient in the GTP bound G105S dimer.

Principal component motion along the first eigen vector for the simulations G105S-FtsZ dimer-GDP (1) and (2) are presented in Fig. 3.6 (c – d). In the simulation G105S-FtsZ dimer-GDP (1) (Fig. 3.6 (c)), the top chain, chain 2, rotates anti-clockwise and the bottom chain rotates from left to right such that the N-terminal domain of the bottom subunit may form dimer interface with the C-terminal domain of the top subunit. The direction of rotation of the respective chains are opposite to what is observed for the simulations G105S-FtsZ dimer-GTP (1) and (2) and similar to the direction of rotation in the simulations WT-FtsZ dimer-GTP (1). This may favour straight protofilament formation in the G105S dimer with GDP.

Principal component motion along the first eigen vector for the simulations G105S-FtsZ dimer-ATP (1) and (2) are presented in Fig. 3.6 (e – f). In the simulation G105S-FtsZ dimer-ATP (1), the top chain, chain 2, rotates anti-clockwise and the bottom chain rotates clockwise. However in the repeat simulation G105S-FtsZ dimer-ATP (2), it is observed that rotation is less and in opposite direction to the respective chains in G105S-FtsZ dimer-ATP (1). It may also be noted that the curvature of the central helix H5 of the respective chains are opposite in the two simulations (Fig. 3.2 (e) and (f)). Since, ATP is not present in the nucleotide binding site of the dimers and forms hydrogen bonds with fewer residues than GTP or GDP, the difference in the direction of rotation of the monomers between the repeat simulations may suggest in the absence of a well bound nucleotide, the chains (protofilament) may take the straight or the curved form.

## Conclusions

We observed that in the G105S monomer simulations, the flexibility of the IDR is greatly reduced in all nucleotide binding states (GTP, GDP and ATP). In our previous studies of the wild-type monomer, we observed that the C-terminal IDR has a much higher flexibility, it makes contacts with HL2-H5 and HC2-SC2. In the presence of a variably bound GTP in the wild-type monomer, it is able to bend the central helix which results in an open conformation of the nucleotide binding site which favours polymerization. The GTP in G105S monomer simulations forms hydrogen bonds with Glu137, Arg141 (and Thr107 in one of the simulations). With the central helix curved towards the nucleotide binding site, it may provide some stabilization to the nucleotide, thus resulting in stable binding. However, since the N-terminal domain is involved in stabilizing the nucleotide, it is not available to form protofilament contacts. Therefore, the G105S bound FtsZ monomer may not be able to polymerize as efficiently as the wild-type monomer. Since, FtsZ GTPase activity is dependent on FtsZ polymerization [3], therefore, in the G105S FtsZ, GTPase activity would be reduced due to its inability to polymerize. In the G105S dimer simulations, it was observed that in the presence of GTP, the bottom chain forms many hydrogen bonds with the nucleotide (8 – 10 hydrogen bonds). It is bound to residues of the signature sequence Ser104, Gly105, Thr107, Thr109 and to S5-H4 loop residues Glu137 and Arg141. The top monomer forms fewer hydrogen bonds, with Glu137 and Arg141. In contrast, in the simulations of the wild-type dimer, the GTP of the dimer interface formed fewer hydrogen bonds, with Arg 141, Thr131 (Asn185 in one of the simulation) and the GTP of the top chain formed hydrogen bonds with Asn23 (in helix H1), Arg141 and Thr131. Our simulations, suggest that the G105S mutation in the signature sequence affects nucleotide interaction. It promotes strong binding with nucleotide in the monomer protein due to which polymerization is not able to occur efficiently. Such type of interaction would result in curved protofilaments with lower subunit contact similar to wild-type FtsZ bound to GDP. Since, the G105S mutant is able to support cell division at permissive temperatures *in vivo*, it is possible that at higher FtsZ concentrations or in the presence of regulatory proteins, protofilaments with wild-type assembly properties are formed. However, in the simulations we observe, that GTP binding does not promote efficient polymerization of G105S FtsZ monomer.

The G105S mutant monomer binds to ATP variably/non-specifically. The flexibility of the C-terminal IDR is reduced. However, due to the variable binding of ATP with the outer residues of the nucleotide binding site, the central helix would still be able to make protofilament contacts. Therefore, FtsZ polymerization may occur (Fig. 3.6 (e), G105S-FtsZ dimer-ATP (1)). However, in the dimer simulations, the nucleotide interactions continue to be non-specific/variable. Therefore, its ATPase activity could not be verified from the simulations.

## Acknowledgement

We would like to thank the Marie Curie Actions program for funding the project, the Minerva supercomputer resource in Warwick University and Iridis supercomputer in Southampton University, Dr. Alison Rodger and Dr. Syma Khalid for supervising the project.

